# Inhibition of the extracellular matrix protein fibulin-3 reduces immunosuppressive signaling in tumor stem cells and increases macrophage activation against glioblastoma

**DOI:** 10.1101/2024.12.28.628031

**Authors:** Somanath Kundu, Soham Mitra, Arivazhagan Roshini, John A. Longo, Sharon L. Longo, Abigail Venskus, Harish Babu, Mariano S. Viapiano

**Affiliations:** Department of Neuroscience and Physiology, State University of New York - Upstate Medical University, Syracuse, NY, USA; Department of Neurosurgery, State University of New York - Upstate Medical University, Syracuse, NY, USA

**Keywords:** brain tumors, immunosuppression, tumor-associated macrophages, extracellular matrix, fibulins, CSF-1, macrophage polarization, NF-kappa B

## Abstract

Glioblastoma tumors remain a formidable challenge for immune-based treatments because of their molecular heterogeneity, poor immunogenicity, and growth in the largely isolated and immunosuppressive neural environment. As the tumor grows, GBM cells change the composition and architecture of the neural extracellular matrix (ECM), affecting the mobility, survival, and function of immune cells such as tumor-associated microglia and infiltrated macrophages (TAMs). We have previously described the unique expression of the ECM protein EFEMP1/fibulin-3 in GBM compared to normal brain and demonstrated that this secreted protein promotes the growth of the GBM stem cell (GSC) population. Here, we describe a novel immunomodulatory role of fibulin-3 and the immuno-boosting effects of targeting this ECM protein. Mice carrying fibulin-3-deficient intracranial tumors showed increased myeloid infiltration and reduced expression of TAM pro-tumoral markers (Arginase, CD206) compared to controls. The opposite was observed in orthotopic tumors overexpressing fibulin-3. *In silico* dataset analysis of clinical datasets revealed positive correlation of fibulin-3 with an immunosuppressive signature, which was validated in GSCs and *in vivo*. We further demonstrated that fibulin-3 regulates the expression of immunosuppressive signals (CSF-1, TGFβ) and the innate immune checkpoint CD47 in GSCs via autocrine activation of NF-κB signaling. Accordingly, immunosuppressive signals were downregulated in GSCs by knockdown of fibulin-3 or inhibition of this protein with an anti-fibulin-3 antibody. Co-culture of GBM cells with syngeneic macrophage lines or primary macrophages in presence of anti-fibulin-3 antibody increased macrophage phagocytosis and antibody-dependent killing of the tumor cells, Furthermore, locoregional delivery of anti-fibulin-3 antibody in mice carrying intracranial GBM increased the infiltration of TAMs expressing pro-inflammatory markers, reducing tumor viability. Our findings show that anti-fibulin-3 approaches, which impact the pericellular ECM surrounding tumor cells and TAMs, can diminish immunosuppression in GBM and boost innate immune responses against the tumor.

---

Advances in cancer immunotherapy have shown unparalleled success against a variety of malignancies and raised hope for improved treatment of malignant gliomas including glioblastoma (GBM), which is the most common and aggressive form of primary brain cancer [1]. However, malignant gliomas remain a formidable challenge for immune-based treatments because of their molecular heterogeneity [2–4], poor immunogenicity [5], and growth in the largely isolated and immunosuppressive neural environment [6, 7].

High-grade gliomas are heavily infiltrated by local microglia and bone marrow-derived macrophages [8, 9], which can make up to a third of total cells recovered from GBM specimens [10, 11]. These innate immune cells are attracted to the tumor mass by the tissue damage and wound-like properties of the tumor [12] that elicit inflammation. Tumor-infiltrating macrophages and microglia (TAMs) are then co-opted by immunosuppressive signals from the tumor, inhibiting further immune responses and unwittingly supporting tumor growth [13, 14]. The resulting “tumor-promoting” transcriptional and phenotypic changes in TAMs have been likened to the M2-like polarized phenotype of macrophages treated with immunosuppressive cytokines [15], although it is well accepted that the phenotypes of TAMs are plastic and heterogeneous and cannot be ascribed to a simple snapshot of gene expression [16]. Nevertheless, it is clear that even different TAM classes have a distinct tumor-promoting role [14] and that, to succeed, GBM immunotherapies must reduce the presence of these cells in the tumor or revert them to an anti-tumor inflammatory status [5]. Remarkably, spatial changes in TAM polarization from the tumor periphery (pro-inflammatory) towards the core (immunosuppressive) correlate with changes in extracellular matrix (ECM) components [17–19], highlighting the potential involvement of the ECM in GBM immunosuppression.

The ECM of malignant gliomas is a complex and unique scaffold surrounding tumor and tumor-associated cells, composed of hyaluronic acid, CNS-specific glycoproteins secreted mainly by astrocytes, and a host of soluble factors and proteins produced by the tumor cells, many of which are absent from the normal brain ECM [20]. Several ECM components are important regulators of the inflammatory or reparative responses of macrophages in GBM [21–23]. For example, secreted hyaluronic acid and the ECM proteins of the tenascin family can limit the infiltration of T cells into GBM [21, 24] and regulate macrophage migration, differentiation, and polarization in these tumors [25, 26]. Nevertheless, the possibility that tumor-secreted ECM molecules can act as inducers of immunosuppression in GBM has remained largely unexplored and therapeutically unexploited.

Fibulin-3 (gene *EFEMP1*) is a pericellular (surface-associated) ECM protein secreted by GBM cells and common to all GBM subtypes but absent from normal brain [27]. Fibulin-3 has multiple tumor-promoting functions in GBM and other solid tumors, such as increasing tumor cell proliferation and invasion [27–30], chemoresistance [31, 32] and the survival of the tumor stem cell population [31, 33]. Fibulin-3 effects can be inhibited with a function-blocking antibody, named mAb428.2, which prevents this protein from activating NF-κB signaling in GBM and PI3K/Akt signaling in mesothelioma, resulting in extended survival of mice carrying orthotopic brain or pleural tumors [28, 34]. Interestingly, this antibody has modest effects against GBM cells cultured alone compared to its more evident apoptotic, anti-angiogenic, and pro-necrotic effects in the tumor mass [34], suggesting that the antibody triggers a stronger than expected anti-tumor response *in vivo*. Indeed, mAb428.2 treatment in mice carrying sub-cutaneous, fibulin-3-expressing, tumors increases the infiltration of macrophages expressing pro-inflammatory cytokines [34], suggesting that targeting of fibulin-3 may increase the response of macrophages against the tumor. However, whether fibulin-3 itself acts as an immunomodulatory factor of GBM TAMs has not been investigated.

Here, we demonstrate that fibulin-3 expression is associated with a significant immunosuppressive signature in GBM and likely contributes to macrophage polarization by an autocrine loop in which fibulin-3 secreted by GBM cells increases their expression of signals such as CSF-1 and CD47, which limit anti-tumor macrophage responses. We further show that targeted knockdown of fibulin-3, as well as inhibition of this protein with a novel, humanized fibulin-3-targeting antibody, reduces the immunosuppressed status of TAMs and increases macrophage cytotoxicity against GBM cells.

## Materials and Methods

### Cells and biological reagents

GBM stem cell (GSC) cultures were established from fresh surgical specimens obtained under informed consent. These cells have been previously described [27, 31, 34, 35] and were re-authenticated using the “Cell Check” service provided by IDEXX-Research Animal Diagnostic Laboratory (Columbia MO). The GBM cell lines U251MG (human), GL261 (mouse) and CNS1 (rat) were cultured as previously described [27, 31, 34, 35]. The pro-monocytic cell lines U937 and THP-1 were cultured in RPMI-1640 (Corning) supplemented with 300 mg/L L-glutamine and 10% fetal bovine serum; these cells were differentiated into macrophages using 100 ng/mL phorbol 12-myristate 13-acetate (Sigma-Aldrich) for 24h (THP-1) or 48h (U937). Human peripheral blood mononuclear cells (PBMCs) were isolated from whole blood (New York Blood Center) and cultured in Iscove’s-modified Dulbecco’s medium (Corning) supplemented with 9% fetal bovine serum and 1% human serum. Mouse myeloid cells were isolated from bone marrow and cultured in RPMI-1640 containing 300 mg/mL glutamine and 10% fetal bovine serum. Human and mouse monocytes were differentiated into mature macrophages using 50 ng/mL M-CSF and 25 ng/mL GM-CSF (Peprotech) for six days. All cultures included standard penicillin/streptomycin antibiotics.

Fibulin-3 cDNA and RNAi sequences have been described and validated previously [27, 31, 34]. Constructs were introduced in GBM cells by lentiviral transduction as described [31]. All GBM cells were transduced with the lentiviral vectors pCDH-EF1-copGFP (System Biosciences) or LPP-hLUC-LV206 (Genecopoeia) for expression of fluorescent or luminescent markers, respectively.

The monoclonal anti-fibulin-3 antibody, mAb428.2, has been described and validated for its ability to inhibit fibulin-3 functions in GBM and other fibulin-3-expressing cancer types [34]. For this study, a fully-humanized variant of mAb428.2 was developed by modifying the original mouse V_H_ and V_L_ sequences and inserting them in a human IgG1 backbone using the Prometheus^TM^ process (Absolute Antibody Ltd) [36]. Humanized mAb428.2 was validated by ELISA, Western blotting, and *in vitro* assays (Suppl. Figure S1), and compared against the predecessor chimeric mAb428.2 (mouse V_H_/V_L_ in a human IgG1 backbone) already reported [28]. Purified reagents using in this study included purified fibulin-3 (R&D Systems), highly-purified human IgG1 and human IgG Fc fragment (Rockland Immunochemicals), and the NF-κB inhibitor caffeic acid phenetyl-ester (CAPE, Cayman Chemical). Commercially available antibodies and PCR primers used in this study are listed in Supplementary Tables I and II, respectively.

### In vitro assays

To measure gene and protein expression, cultured cells were dissociated mechanically, without trypsin treatment, and processed for semiquantitative RT-PCR or Western blotting following standard procedures. RNA was collected from transfected cells using Pure Link RNA mini kit (Invitrogen). Lipofectamine-mediated transfections for transient gene overexpression or knock-down were performed 24-48 hours before cell collection. Macrophages, differentiated from cell lines or primary cells, were co-cultured with GBM cells in 96-well plates at an effector: target: ratio of 5:1 (5×10^3^ macrophages/well) for 48 hours to assess GBM cytotoxicity, and at a E:T ratio of 1:1 (5×10^5^ macrophages/well) for 60 minutes to visualize phagocytic activity. Cytotoxicity was quantified by a decrease in fLuc signal of the tumor cells (BrightGlo Luc kit, Promega) and further validated in some experiments by measuring released LDH enzyme as a separate measure of cell death (CytotTox kit, Promega). To quantify phagocytosis of GFP-expressing GSCs, co-cultures were lightly fixed with 4% phosphate-buffered paraformaldehyde and prepared for flow cytometry following standard procedures, using CD45 to gate the macrophage population. Alternatively, following cell fixation, some co-cultures were processed for confocal microscopy to visualize GSC-derived fragments inside the macrophages.

### In vivo studies

All animal experiments were performed following the institutional IACUC approval at SUNY Upstate Medical University. GSCs (2 µL containing 1×10^4^ cells/µL) were implanted in the right-side striatum of athymic nude mice (FoxN1^nu/nu^, 1:1 female:male ratio, The Jackson Laboratory) as previously described [31]. GL261 (2 µL containing 5×10^4^ cells/µL) and CNS1 cells (3 µL containing 5×10^4^ cells/µL) were implanted in the same brain region of C57Bl/6 mice and Lewis rats [27], respectively. To induce expression of fibulin-3 shRNAs mice received doxycycline (1 mg/mL) in drinking water containing 2 % w/v sucrose, and were allowed to drink medicated water *ad libitum* after recovering from surgery. Fibulin-3 cDNA was overexpressed constitutively and needed no induction. Tumors were allowed to grow for 14 to 21 days, after which the animals were euthanized, perfused with phosphate buffered saline and paraformaldehyde, and their brains processed for immunohistochemistry. In parallel, some brains were freshly resected, dissociated mechanically and enzymatically (1 mg/mL type-IV collagenase and 0.25 mg/mL DNAse I, Worthington Biochemical) and fractionated in a Percoll gradient (Cytiva). The enriched mononuclear immune cell fraction was processed for semiquantitative RT-PCR or by flow cytometry using a LSRFortessa cell analyzer (Becton Dickinson). Cytometry results were analyzed using FlowJo v.10.1.

To treat intracranial tumors with humanized anti-fibulin-3 Ab428.2 or control IgG1 antibodies, animals were operated a week after tumor injection to implant an intracranial canula and a sub-cutaneous osmotic reservoir (Alzet #2001 containing 5 mg/mL antibody) as described [34]. Animals received antibody infusion for 8 days and were euthanized 48 hours after completing the antibody treatment.

### External datasets and data analysis

Expression of fibulin-3 and immunomodulatory genes was extracted from the IDH1-wildtype GBM datasets available at the Cancer Genome Atlas and Chinese Glioma Genome Atlas databases, accessed using the aggregator website Gliovis [37]. The bulk transcriptional subtypes (proneural, mesenchymal, classical) of GBM were identified using the approach proposed by Wang et. al. [38]. NF-κB-regulated genes were compiled from datasets listed at http://bioinfo.univ-lille.fr and http://www.bu.edu/nf-kb/gene-resources/target-genes [39, 40].

To perform RNA sequencing of transfected cells, libraries were prepared from poly-A-enriched RNA and sequenced in an Illumina HiSeq instrument with 2×150bp paired-end reads and an average coverage depth of >25M reads/sample. Sequencing reads were mapped to the hg38 human genome version and quantified using Partek Flow software. Gene counts were processed to remove cell line-dependent batch effects and quantified as trimmed mean of M-values (TMM) for analysis. Gene ontology was analyzed using Gene Set Enrichment Analysis (GSEA) tools provided by Partek Flow. Raw and processed gene counts have been deposited in the NCBI Gene Expression Omnibus (GSE284269).

All *in vitro* experiments were repeated in triplicate with three independent replicates per condition. *In vivo* studies used N=5-8 per condition. Immunohistochemical quantification was performed by investigators blinded to the experimental conditions, using at least 5 separate tissue sections per animal. Image analysis used ad-hoc scripts in ImageJ (v. 1.54) to quantify IBA1-positive, CD206-positive, and co-localized IBA1/CD206 pixels in each tumor. Values in the graphs represent mean ± SD. Comparisons between two conditions were analyzed by Student’s t-test or multiple t-tests with multiple comparisons’ correction. Grouped results were analyzed by one- or two-way ANOVA depending on the experimental design. All differences were deemed significant at p<0.05. Statistical differences in figure panels are represented as *<0.05; **<0.01; and ***<0.001.

## Results

### Fibulin-3 expression by GBM cells correlates with the infiltration of tumor-promoting macrophages

Fibulin-3 is known to have multiple tumor-promoting effects in GBM, such as enhancing tumor cell invasion [27], stemness [31, 32], vascularization [35], and peritumoral astrocytosis [33]. However, whether this protein affects TAMs infiltrating the tumor has remained unexplored. To address this question, we started by analyzing macrophage infiltration in control and fibulin-3-deficient proneural GSC-derived tumors (GBM08 cells) implanted in athymic mice compatible with these xenografts. Our results showed a significant increase of infiltrated, IBA1-positive macrophages in fibulin-3-deficient tumors compared to controls (Figure 1A-B). At the same time, we saw a significant decrease in the expression of CD206 in those macrophages, which is characteristic of the M2-like, tumor-promoting phenotype of TAMs. These results were confirmed with a second, more aggressive, mesenchymal GSC model (GBM34 cells, Suppl. Figure S2A). Importantly, to avoid quantification bias due to the intrinsic negative effects of fibulin-3 knockdown on the tumor cells, we chose tissue sections of similar cross-sectional area and cellular density in all animals (quantified in Suppl. Figure S2A). Athymic mice are considered to retain a functional macrophage population able to deliver anti-tumor inflammation [41–43]; nonetheless, we further validated our observations in an immunocompetent model using control and fibulin-3-deficient CNS1 cells implanted in Lewis rats (Suppl. Figure S2A). Our results in CNS1-derived tumors were essentially the same as observed with GSCs.

**Figure 1:**
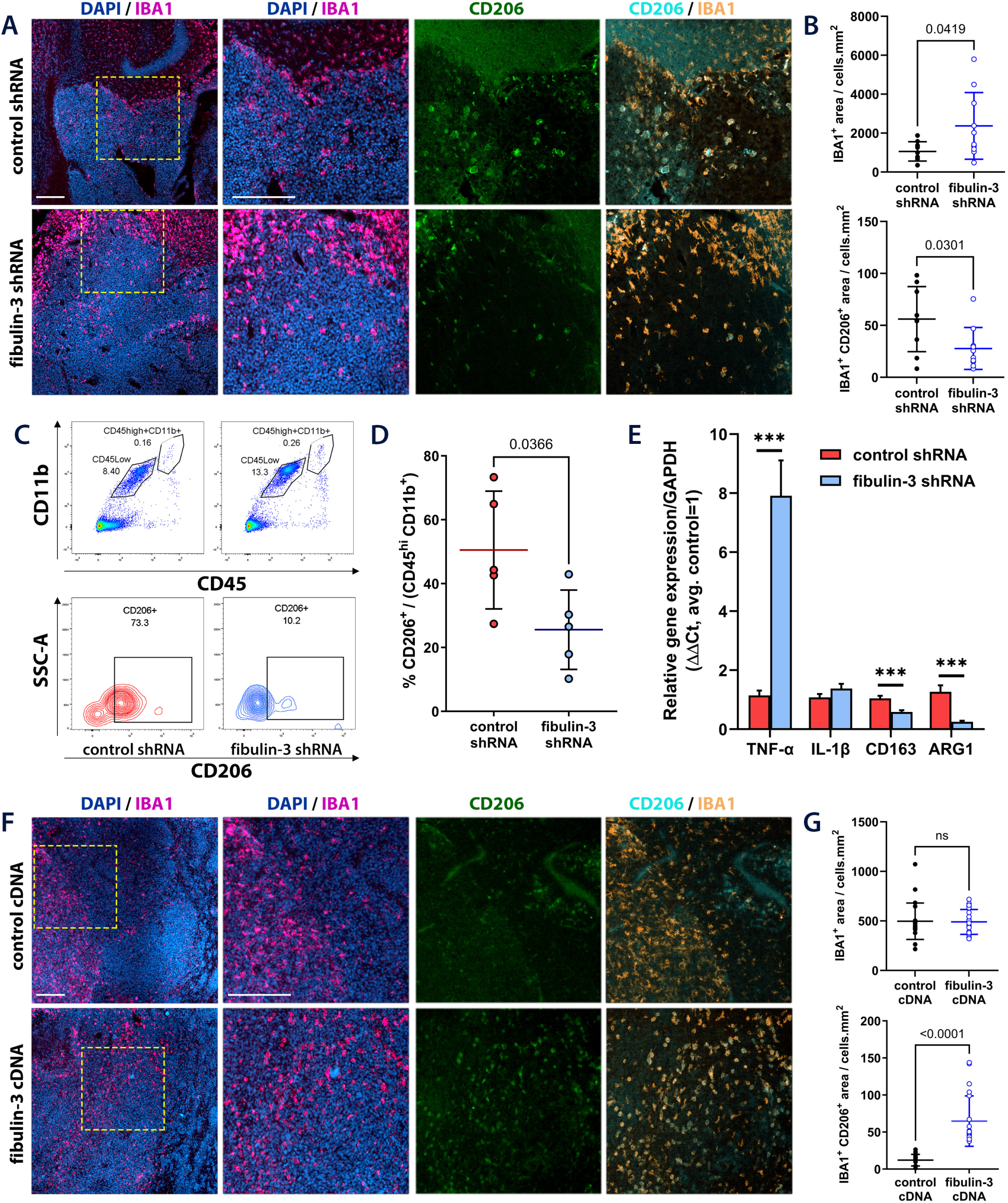
Fibulin-3 expression affects the phenotype of tumor-infiltrating macrophages. **A)** Representative images of GSC-derived intracranial tumors (GBM08 line) expressing control or fibulin-3 shRNAs, showing TAM infiltration (IBA1 staining) and expression of the immunosuppressive marker CD206 in TAMs; cell nuclei were stained with DAPI (scale bars: 200 µm). **B)** Quantification of TAM infiltration (IBA1^+^ area within the tumor) and IBA1/CD206 co-expression in the infiltrated TAMs. Results analyzed by Welch’s-corrected t-test. **C)** Representative flow cytometry gating of the TAM populations (CD45^hi^ CD11b^+^ and CD45^lo^ CD11b^+^) recovered from the tumors indicated in (**A**). **D)** Flow cytometry quantification of CD206+ TAMs recovered from freshly resected control and fibulin-3-deficient tumors; comparison by Student’s t-test. **E)** qRT-PCR for selected genes expressed in TAMs recovered from control and fibulin-3-deficient tumors; analysis by multiple paired t-tests with correction for multiple comparisons. **F)** Representative images of CNS1-derived intracranial tumors expressing control or fibulin-3 cDNA; stains are the same as in (**A**); scale bars: 200 µm. **G)** Quantitative analysis of the tumors in (**F**) showing TAM infiltration (IBA1^+^ area within the tumor) and IBA1/CD206 co-expression in the infiltrated TAMs. Results analyzed by Welch’s-corrected t-test.

To confirm our quantitative immunohistochemistry results, we isolated TAMs from freshly resected control and fibulin-3-deficient GBM34 tumors and quantified CD206 expression in the CD45^+^/CD11b^+^ population, observing a significant decrease of CD206 in TAMs isolated from fibulin-3-deficient tumors (Figure 1C-D). Furthermore, we profiled those TAMs by qRT-PCR and observed a significant increase in the mRNA expression of a pro-inflammatory marker (TFN-α) with simultaneous decrease of genes associated with immunosuppressed phenotype (CD163 and Arginase-1, Figure 1E).

Next, we performed a similar immunohistochemical quantification of tumors overexpressing fibulin-3. We chose two cell models (CNS1 and GL261) that have modest expression of fibulin-3 and have been validated for overexpression of this protein before [27]. Results with CNS1 cells implanted in Lewis rats did not show significant changes in total macrophage infiltration (IBA1-positive cells) but demonstrated significant increase of CD206 expression in those TAMs (Figure 1F-G). Results with GL261 cells implanted in C57Bl/6 mice (Suppl. Figure S2B) showed a modest decrease in TAM infiltration and large increase in their CD206 expression.

Taken together, these results strongly suggest that high expression of fibulin-3 by GBM cells correlates with immunosuppressed (i.e. CD206 positive) features of infiltrated TAMs, whereas decreased expression of fibulin-3 correlates with increases infiltration of TAMs having a less immunosuppressed and more inflammatory-prone profile. Accordingly, we next investigated whether fibulin-3 could be considered an immunosuppressive factor in GBM.

### Fibulin-3 correlates with an immunosuppressive signature in GBM tumors and GSCs

Fibulin-3 activates NF-κB signaling in GBM cells and, accordingly, correlates with the expression of numerous NF-κB-regulated genes [33]. Several immunosuppressive signals have been identified as direct targets of NF-κB regulation at the transcriptional level (see compiled list at http://www.bu.edu/nf-kb/gene-resources/target-genes), including the cytokines CSF-1, TGFβ, and IL-10, as well as the checkpoint inhibitors CD274 (PD-L1), CD80, and CD47, all of which form an immunosuppresive signature. We analyzed the expression of these genes in clinical GBM specimens using TCGA and CGGA datasets and found a significant positive correlation of fibulin-3 with this immunosuppressive signature (Figure 2A and Suppl. Figure S3). Expectedly, the cumulative correlation of fibulin-3 with immunosuppressive signals was highest in mesenchymal tumors (Figure 2B), which have the highest expression of fibulin-3 [31] and predominant immune component [38].

**Figure 2:**
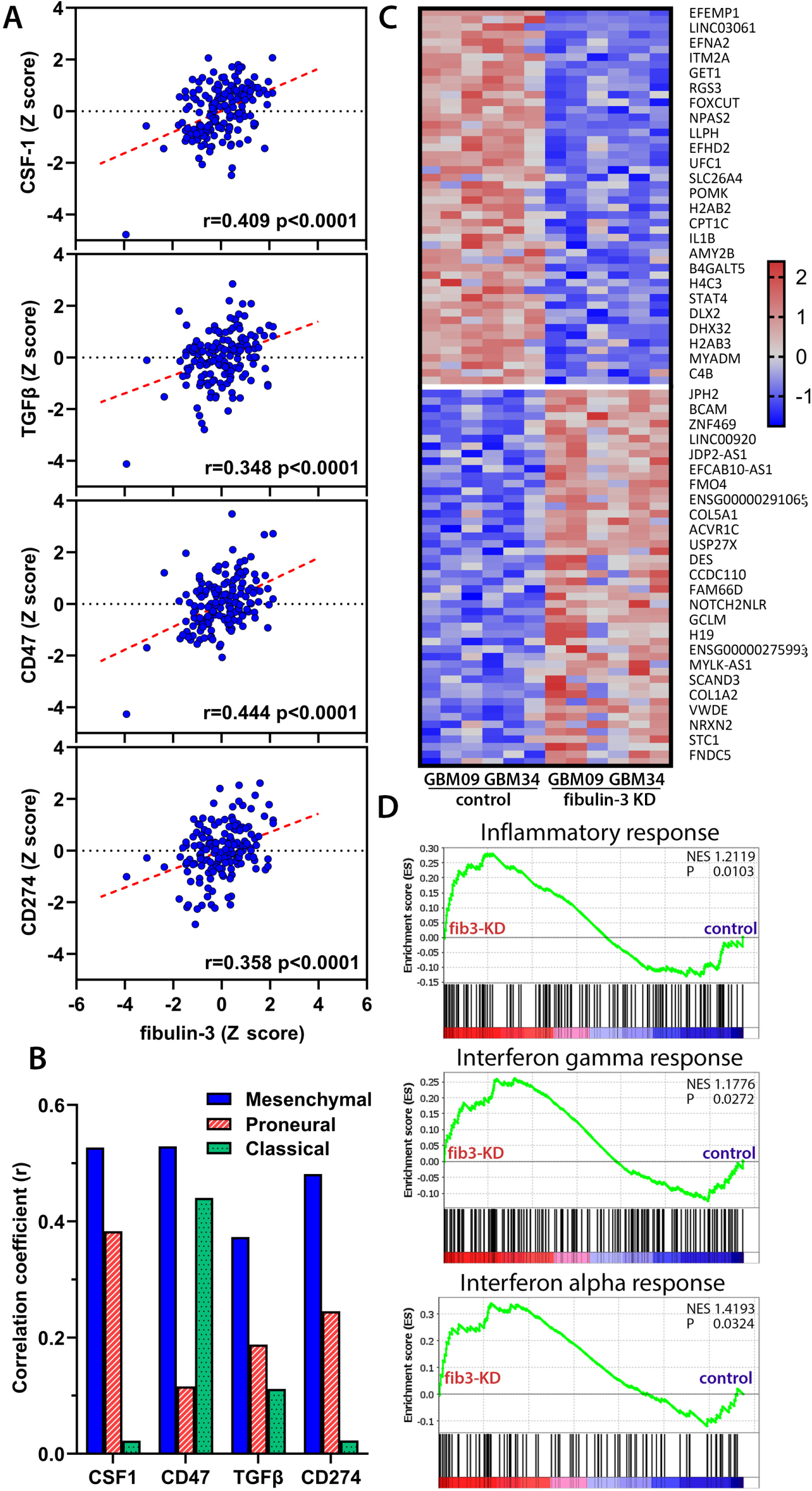
Fibulin-3 expression correlates with an immunosuppressive signature in GBM. **A)** Correlation of fibulin-3 mRNA expression with the expression of CSF-1, TGFβ, and the checkpoints CD47 and CD274 (PD-L1), extracted from the TCGA GBM dataset. **B)** Correlation of fibulin-3 co-expressed with the markers indicated in (**A**) in GBM tumors separated by their bulk transcriptional subtype. **C)** Heatmap showing top gene expression changes in two GSC lines (GBM34, GBM09) stably transfected with control or fibulin-3 shRNAs. **D)** Changes in gene expression observed in hallmark signatures for inflammatory response and signaling by interferon gamma and alpha (data analyzed using GSEA v.4.3.3), comparing fibulin-3-deficient GSCs (*fib-3 KD*) versus control cells.

We validated these findings by knocking-down fibulin-3 in two mesenchymal GSC models with high endogenous fibulin-3 expression (GBM09 and GBM34 models) and analyzed the resulting transcriptomic changes in these cells (Figure 2C). Gene ontology analysis revealed increased expression of genes associated with inflammatory, interferon-gamma, and interferon-alpha responses (Figure 2D) after knockdown of fibulin-3, suggesting that fibulin-3 may downregulate these genes in GSCs.

To further confirm our results suggesting this immunosuppressive function of fibulin-3 in GSCs, we validated immunosuppressive genes expressed in GSCs with direct impact on macrophage phenotype. We focused on the expression of the cytokines CSF-1 and TGFβ, as well as the cell-surface receptor CD47 that inhibits phagocytosis and acts as innate immune checkpoint. In all cases, knockdown of fibulin-3 in GSCs decreased the expression of these factors at the mRNA and protein levels (Figure 3A-B). We returned to our intracranial GSC-derived fibulin-3-deficient tumors (analyzed in Figure 1A-E) and performed qRT-PCR for human-specific transcripts. In agreement with our prior results, we observed that fibulin-3-deficient tumors had a much lower expression of multiple immunosuppressive signals compared to control tumors (Figure 3C), which would explain the increased infiltration of macrophages and their decreased CD206 expression. Finally, we transiently overexpressed fibulin-3 cDNA in GBM cell lines (U251MG, GL261) as well as a colorectal cancer cell line that lacks endogenous fibulin-3 expression (Colo201 [34]). In all these cases, the forced overexpression of fibulin-3 significantly increased the expression of CSF-1 and CD47 that suppress anti-tumor innate immunity (Figure 3D).

**Figure 3:**
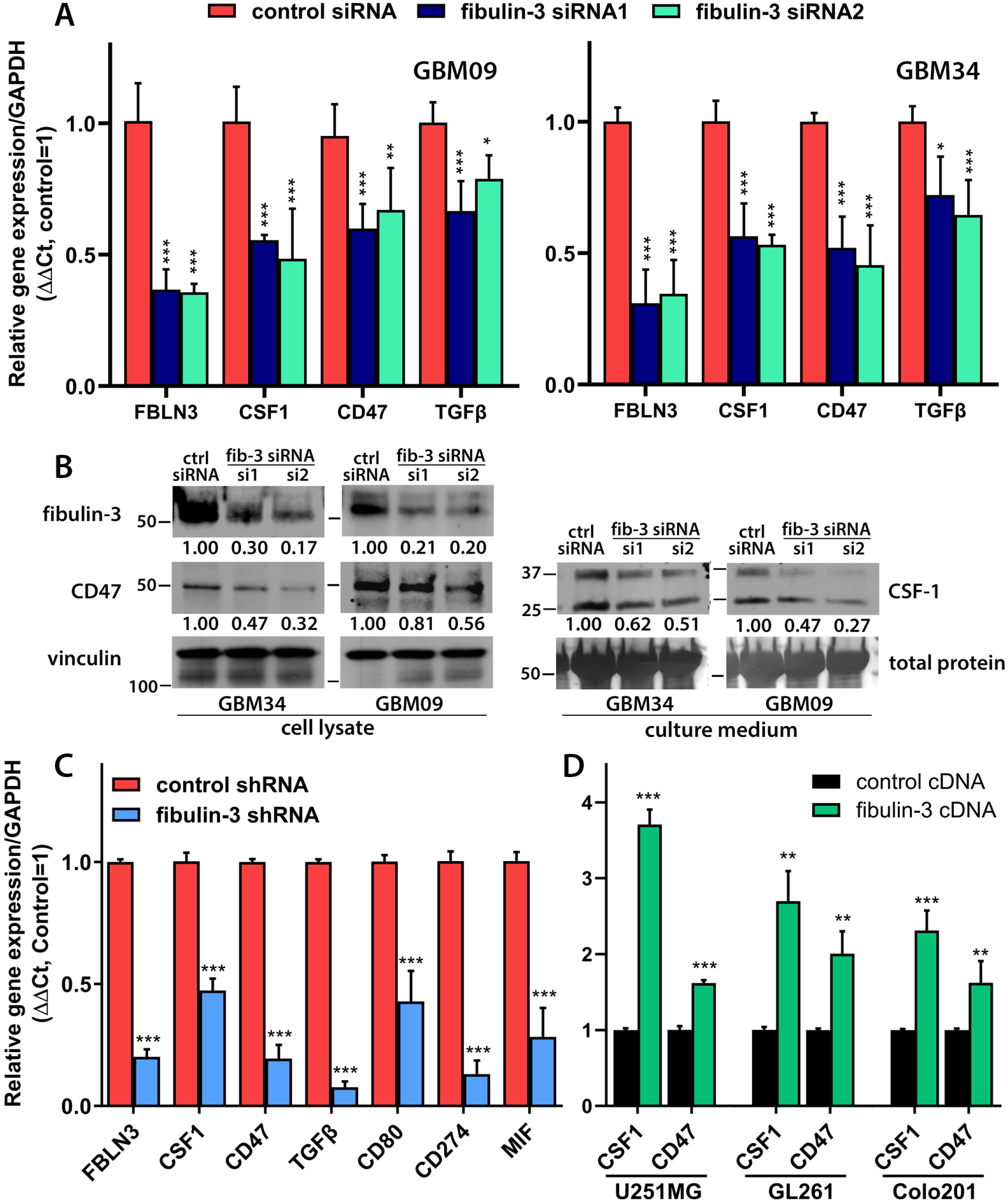
Fibulin-3 regulates the expression of immunosuppressive signals in GBM cells. **A)** Transient transfection of fibulin-3 siRNAs in GSCs results in decreased expression of the cytokines CSF-1 and TGFβ and the innate immune checkpoint CD47, all of which modulate TAM behavior. **B)** Same results as in (**A**), validated by Western blotting. Total protein loading (amido black stain of blotting membranes) was used as loading control to normalize the expression of proteins detected in culture medium. The numbers indicated relative integrated optical density for each band, normalized to the loading control and to total protein measurement. **C)** qRT-PCR of human fibulin-3 (*FBLN3*) and immunosuppressive signals detected in intracranial GSC-derived tumors expressing control or fibulin-3 shRNAs (human-specific primers were used to detect human mRNA in tissue resected from mouse brains). Statistical analysis performed by multiple paired t-tests with multiple-comparisons’ correction. **D)** Expression of CSF-1 and CD47 in the GBM cell lines U251MG and GL261, transfected to overexpress fibulin-3. Experiments were repeated in the carcinoma cell line Colo201 that lacks endogenous fibulin-3 expression. Statistical analysis by one-way ANOVA for each cell line.

### Fibulin-3 drives the expression of immunosuppressive signals in GBM cells via NF-κB signaling

The results above strongly suggest that fibulin-3 induces the expression of an immunosuppressive signature in GBM cells that can affect the phenotype of tumor-infiltrating macrophages. We further explored the mechanism by which fibulin-3 may regulate macrophage immunosuppressive signals (CSF-1, CD47) in GBM. We investigated this in a conventional cell line (U251MG) that is efficiently transfected with fibulin-3 cDNA, as well as a mesenchymal GSC line (GBM09) that was directly exposed to purified fibulin-3 (500 ng/mL).

Transfection of U251MG cells with fibulin-3 cDNA resulted in significantly increased expression of CSF-1 mRNA as expected (Figure 4A-B). This effect was largely diminished by knockdown of the canonical NF-κB transcription factor P65/RelA (following a previously validated procedure [33]) and abolished by treating the cells with the low-toxicity NF-κB inhibitor, CAPE (20 µM). NF-κB inhibition with CAPE also inhibited completely the increased expression of CD47 by fibulin-3 in U251MG cells (Figure 4C). Similar results were observed in GBM09 cells: The enhancing effect of purified fibulin-3 on the expression of CSF-1 and CD47 was significantly diminished by knockdown of P65/RelA (Figure 4D) or CAPE treatment (Figure 4E-F). Taken together with our prior results, this confirmed that fibulin-3 drives the expression of an immunosuppressive signature in GBM cells via NF-κB activation.

**Figure 4:**
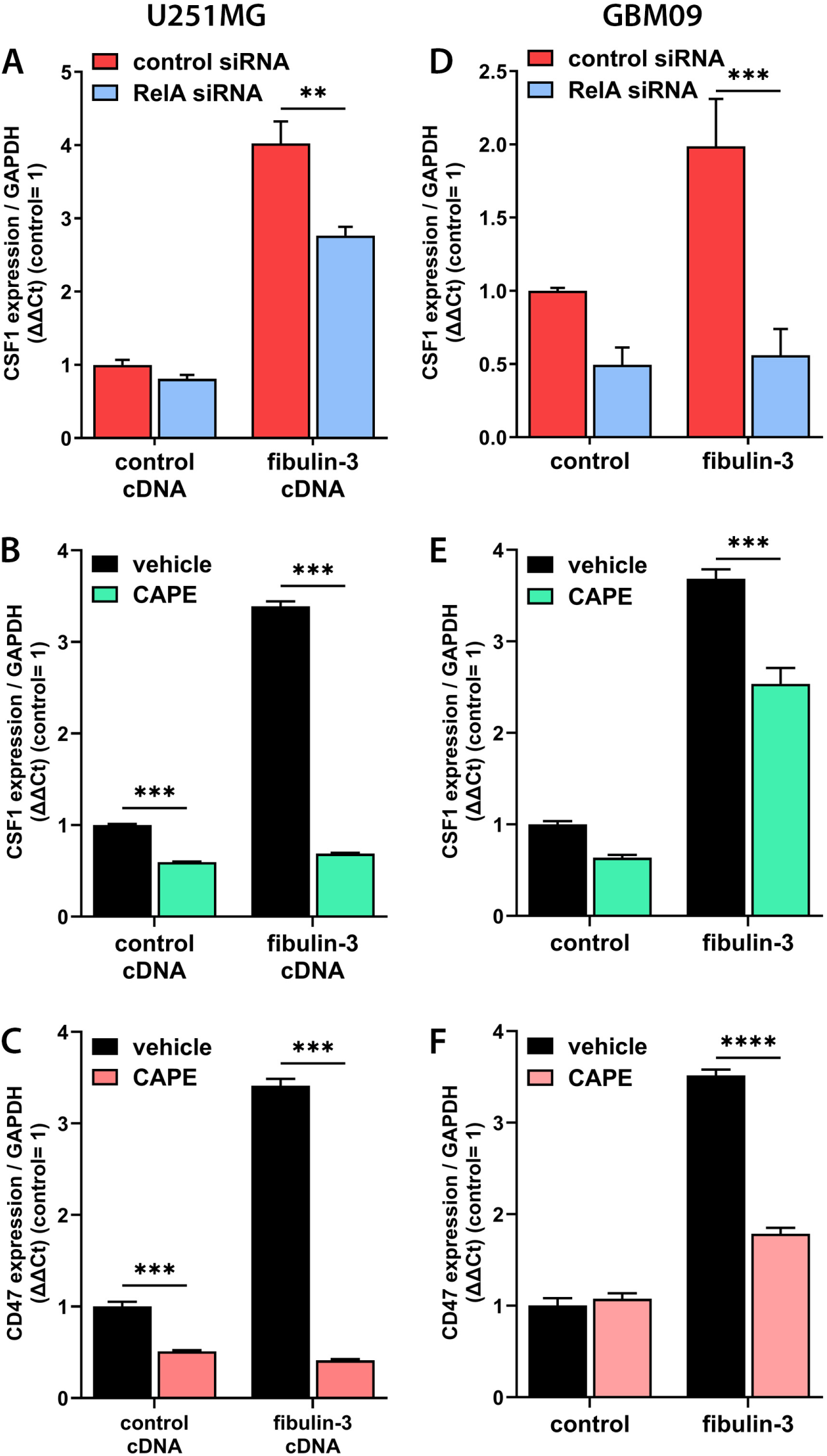
Fibulin-3 regulates the expression of immunomodulatory signals in GBM cells via NF-kB signaling. **A)** Expression of CSF-1 mRNA in U251MG cells co-transfected with fibulin-3 cDNA and p65/RelA siRNA, or their respective controls. **B-C)** Expression of CSF-1 (**B**) and CD47 (**C**) mRNAs in control and fibulin-3-overexpressing U251MG cells in presence of the NF-kB inhibitor, CAPE (20 µM), or its vehicle. **D)** Expression of CSF-1 mRNA in GBM9 GSCs transfected with control or p65/RelA siRNAs and exposed overnight to purified fibulin-3 (500 ng/mL). **E-F)** Expression of CSF-1 (**E**) and CD47 (**F**) mRNAs in GBM9 cells exposed to purified fibulin-3, CAPE, or their respective vehicles. Results in all panels analyzed by two-way ANOVA.

### Blockade of fibulin-3 increases infiltration of inflammatory macrophages that target GBM cells

We next investigated if inhibition of fibulin-3 after tumor formation could be sufficient to prevent the immunosuppressed phenotype of TAMs *in vivo*. To this end, we implanted intracranial tumors (mesenchymal GBM09 cells) in athymic mice and, once the tumors were established, we locally infused humanized anti-fibulin-3 mAb428.2 [34] or a control IgG1. Tumors were processed shortly after completing the antibody infusion and analyzed by quantitative immunohistochemistry for IBA1 and co-localized IBA1/CD206 as shown in Figure 1 (results were again normalized to tumor cross-sectional area and total cell density, Suppl. Figure S2C).

Tumor treatment with mAb428.2 resulted in a significantly increased infiltration of macrophages that had a lower expression of the immunosuppressive marker, CD206, compared to controls (Figure 5A-C). In agreement, qRT-PCR of resected tumor tissue using mouse-specific primers demonstrated a significant decrease of immunosuppressive markers and a trend towards upregulation of pro-inflammatory markers in the tumors treated with anti-fibulin-3 antibody (Suppl. Figure S4). Together, these results closely matched our prior results in which fibulin-3 had been knocked down (Figure 1), suggesting that the antibody was achieving a comparable effect. In agreement, treatment of different fibulin-3-expressing, mesenchymal GSC lines with mAb428.2 (50 µg/mL) significantly decreased their expression of CSF-1, TGFβ, and CD47 (Figure 5D).

**Figure 5:**
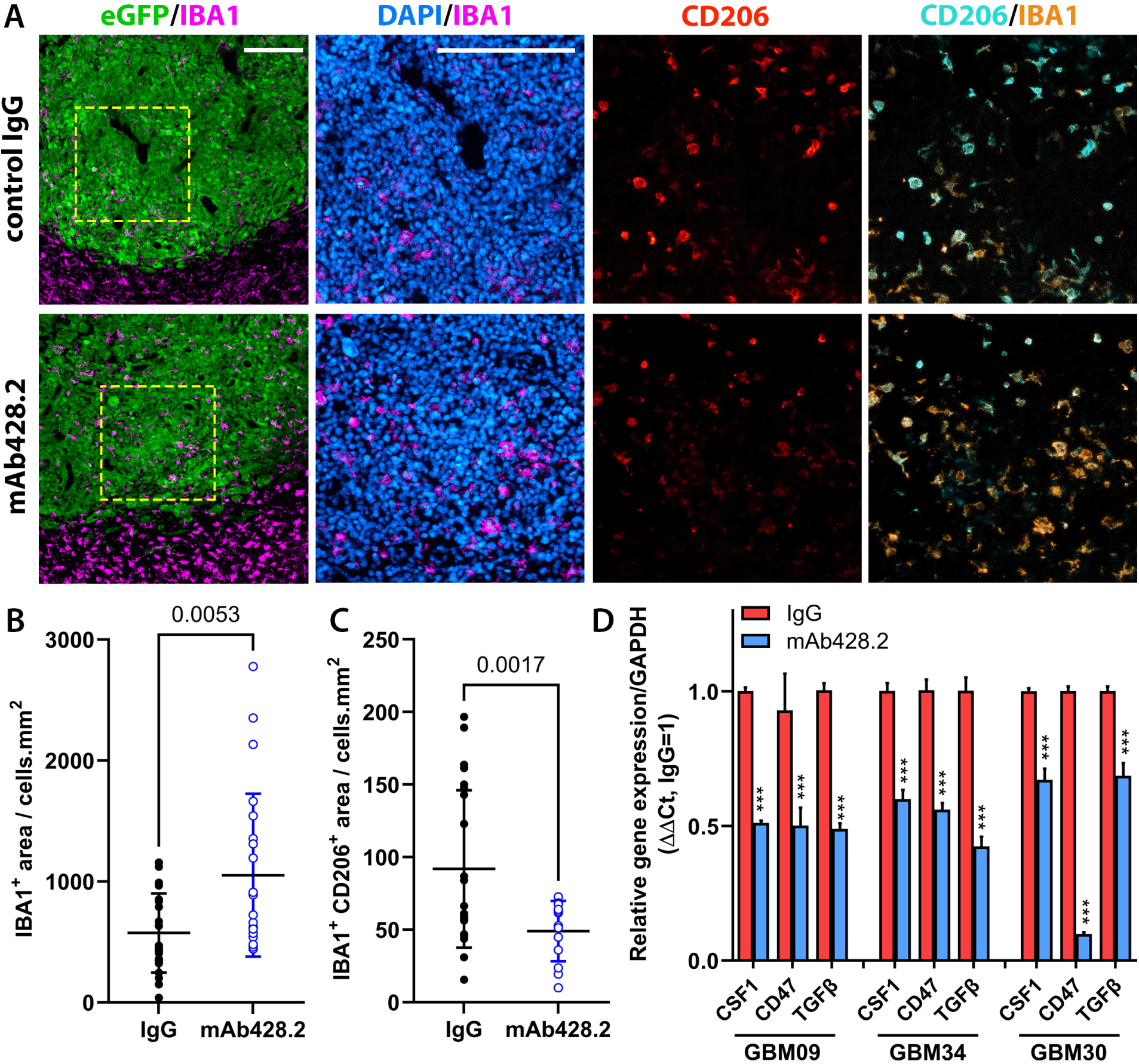
Anti-fibulin-3 antibody increases infiltration of pro-inflammatory macrophages in GBM. **A)** Representative images of intracranial GSC-derived tumors (GBM09) treated with humanized anti-fibulin-3 mAb428.2 or its IgG control. Tissues were stained to reveal TAM infiltration (IBA1) and expression of CD206 in infiltrated TAMs (scale bars: 200 µm). **B-C)** Quantification of TAM infiltration (**B**) and CD206 expression in infiltrated TAMs (**C**) in tumors treated with mAb428.2 or its control; data analyzed by Welch’s corrected t-test. **D)** Expression of CSF-1, TGFβ, and CD47 in three different GSC cultures treated with mAb428.2 or its isotype control. Analysis by multiple paired t-tests for each GSC line.

We next tested if mAb428.2 could modify the behavior of macrophages co-cultured with GBM cells. Using the monocytic cell line U937, differentiated to macrophage phenotype, we established co-cultures of macrophages and GBM cells (either GSCs or U251MG cells). We monitored cytotoxicity of the tumor cells alone or co-cultured with macrophages in presence of mAb428.2 or in different control conditions. In all cases, GBM cell death was significantly increased when tumor cells were co-cultured with macrophages in presence of mAb428.2 compared to IgG1 or vehicle controls (Figure 6A). These effects were validated using a different macrophage cell line (THP-1 cells), as well as primary macrophages derived from human PBMCs or from mouse bone marrow (Suppl. Figure S5). In all cases, co-culture of GBM cells with their syngeneic macrophages resulted in higher cytotoxicity of the tumor cells when mAb428.2 was added to the -cultures.

**Figure 6:**
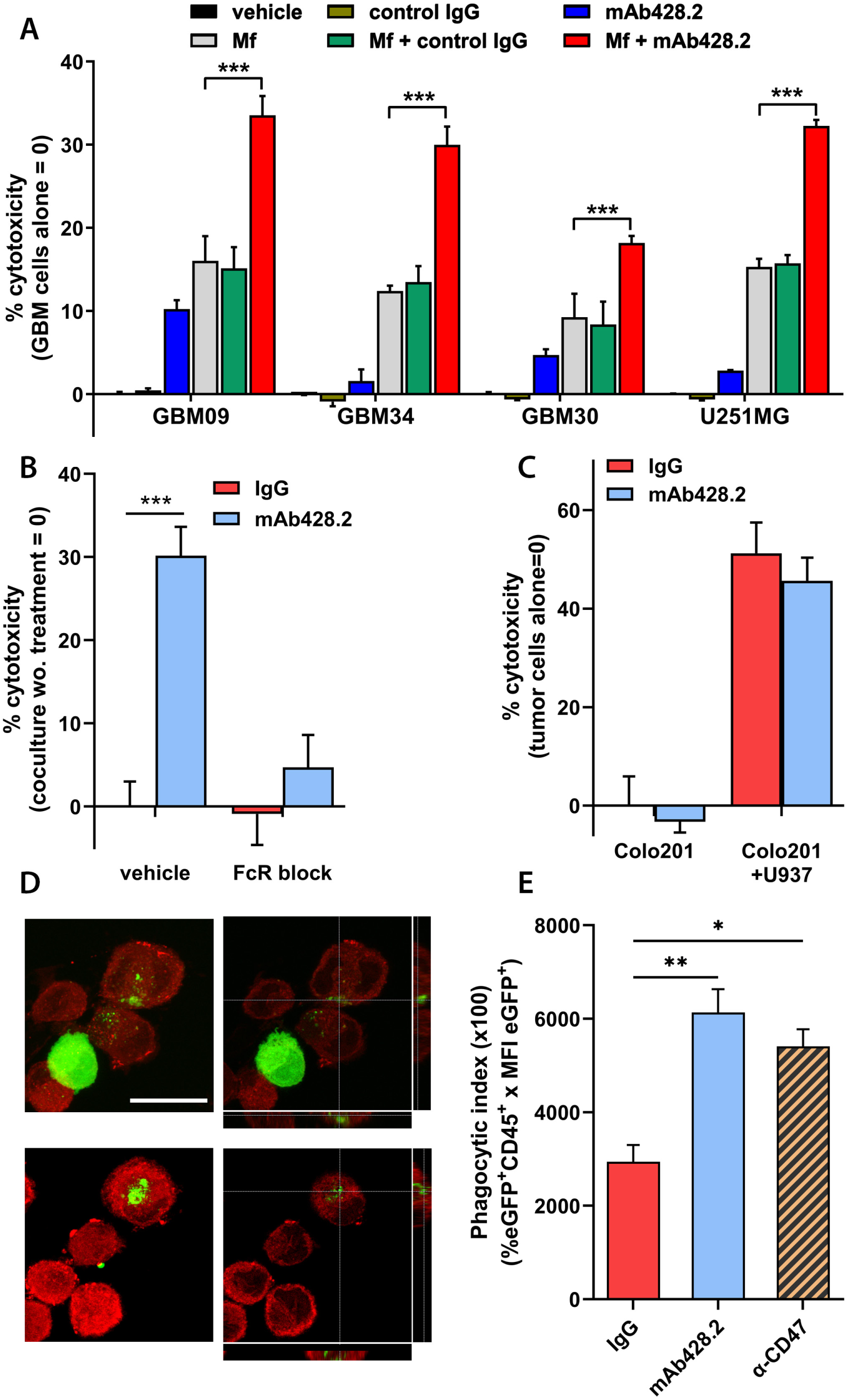
Anti-fibulin-3 increases macrophage attack of GBM cells. **A)** Quantification of cytotoxicity in fLuc-expressing GBM cells co-incubated with U937 macrophages (E:T = 5:1) in presence of anti-fibulin-3 mAb428.2, control IgG, or vehicle. Control conditions included GBM cells incubated alone in presence and absence of the same antibodies. Results analyzed by one-way ANOVA for each cell line. **B)** mAb428.2-induced cytotoxicity of GBM9 cells co-cultured with U937 macrophages that were not pretreated (*vehicle*) or pre-treated with excess of IgG-Fc fragment to block macrophage Fc receptors (*FcR block*). **C)** mAb428.2-induced cytotoxicity of Colo201 cells cultured alone or with U937 macrophages, in presence of mAb428.2 or its control IgG. Results in (**B**) and (**C**) analyzed by two-way ANOVA. **D)** Representative images of DiI-stained macrophages (red) co-cultured with eGFP-expressing GBM9 cells (green) in presence of mAb428.2. Confocal sectioning was performed to confirm that eGFP particles were present inside the macrophages; scale bar: 25 µm. **E)** Flow cytometric analysis of phagocytic activity of U937 macrophages co-cultured with GBM9:eGFP cells in presence of control IgG, mAb428.2 (50 µg/mL), or an antibody against the cell-surface phagocytosis inhibitor CD47 (10 µg/mL, positive control). Results analyzed by one-way ANOVA.

To validate the specificity of this effect, we repeated co-cultures of GBM cells with macrophages that had been pre-treated with an excess of purified human IgG Fc fragment (100 µg/mL) to saturate Fc receptors [44]. This pre-treatment prevented the death of the tumor cells in co-culture (Figure 6B), suggesting that the mechanism of death involved macrophages attacking mAb428.2-opsonized tumor cells. Separately, we evaluated mAb428.2 in co-cultures of macrophages with the cancer cell line Colo201 that does not express fibulin-3. Although Colo201 had higher baseline cytotoxicity in co-culture compared to GBM cells (likely due to the low expression of the inhibitor CD47 in Colo201), the death of these cells was unaffected by mAb428.2 (Figure 6C). This suggested that the antibody-dependent cell cytotoxicity of macrophages against GBM cells was specific to recognition of pericellular fibulin-3. We recovered macrophages co-cultured with eGFP-expressing GSCs in presence of mAb428.2 and confirmed by confocal microscopy the presence of eGFP-positive particles inside the macrophages (Figure 6D), suggesting that the antibody had increased immune attack by phagocytosis/trogocytosis. Finally, co-cultures of macrophages with eGFP-expressing GSCs were analyzed by flow cytometry and a phagocytic index was defined as the median fluorescence intensity of eGFP detected in the macrophage (CD45^+^) population. Addition of mAb428.2 significantly increased the phagocytic index in co-cultures compared to control IgG (Figure 6E).

## Discussion

Malignant gliomas contain a complex extracellular matrix whose components are produced both by the tumor cells and by a sizable population of tumor-associated neural cells. GBM ECM molecules include glycoproteins and proteoglycans that are highly specific to neural tissue [45] as well as fibrillar and fibril-associated proteins absent from normal brain and produced by tumor cells undergoing a mesenchymal-like differentiation program [46], or by tumor-associated pericytes [47]. The result is a unique tissue scaffold that is different from the mucinous ECM composition of normal brain parenchyma and from the fibrillar ECM scaffold of other solid tumors. It is therefore not surprising that ECM proteins produced “*de novo*” by GBM cells have significant impact on processes such as tumor cell invasion, tumor neo-vascularization, astrocytic reactivity, and regulation of immune cell responses in the neural microenvironment.

Anti-ECM approaches have demonstrated significant effect against GBM, either by direct effect against the tumor cells [34, 48] or as adjuvant treatment to improve penetration of oncolytic viruses [49] and immune cells [24]. Antibodies against ECM proteins have successfully reached clinical trials for GBM, such as *tenatumomab* against tenascin-C [50, 51] and *L19*, an antibody against a tumor-specific variant of fibronectin [52–54]. It is worth noting that most antibodies against ECM components do not directly block the functions of these proteins and therefore depend on conjugated payloads (e.g., 131-Iodine or IL-2) to achieve anti-tumor effect. This does not apply to anti-fibulin-3 mAb428.2, which was specifically designed to bind and inhibit the signaling motif of its target [34].

In the present investigation we have shown that the pericellular protein fibulin-3, which is part of the glycocalyx and pericellular ECM, has a significant immunomodulatory role that limits the ability of TAMs to act against GBM cells. This effect is exerted by an autocrine loop in which fibulin-3, secreted by GBM cells, activates NF-κB in these cells to increase the expression of immunosuppressive factors such as CSF-1, TGFβ, and the phagocytosis inhibitor, CD47. This is, to our knowledge, the first observation of a “small fibulin” member (i.e., fibulin-3, -4, and -5) as a potential immunomodulatory factor. The ability of fibulin-3 to indirectly modify the behavior of TAMs is particularly interesting because of the spatial localization of this protein in GBM tumors. Fibulin-3 is enriched in peri-necrotic areas of the tumor core, accumulating around blood vessels that are the route of entry of bone-marrow sourced macrophages [31, 35]. Tumor areas rich in fibulin-3 would therefore be primed to induce macrophage immunosuppression upon infiltration, contributing to the rapid hijacking of this immune population.

The upregulation of multiple macrophage-suppressive signals by fibulin-3, such as CSF-1 and TGFβ, is also of particular interest because it suggests that multiple, redundant, mechanisms converge to promote the TAM tumor-promoting phenotype [9, 14] that has been highlighted as a major contributor to the limited success of GBM immunotherapy to date [5]. In this regard, we noted an interesting aspect of CSF-1 being downregulated by fibulin-3 inhibition, contrasting with the seminal work of Joyce and collaborators that demonstrated the powerful polarizing effect of this cytokine on GBM TAMs [55, 56]. Sustained inhibition of CSF-1 receptor in TAMs, shown by Quail et al., results in upregulation of IGF-1 in these cells, which leads to a rebound of the tumor via IGF-1/IGF1R pro-tumorigenic signaling [57]. In contrast, downregulation of CSF-1 in GSCs by fibulin-3 inhibition did not lead to IGF-1 upregulation in the tumor (Suppl. Figure S4), possibly preventing this tumor-salvage pathway. Whether this reveals different effects of inhibiting CSF-1R compared to decreasing CSF-1 expression, or whether the lack of IGF-1 upregulation was caused by other factors – such as the absence of TAM-regulating T cells in the athymic model – remains to be investigated.

Another interesting effect of fibulin-3 inhibition is the downregulation of the checkpoint CD47 in tumor cells, which prevents phagocytosis by engaging the macrophage receptor SIRP-1α [58]. CD47 expression has been correlated with poor prognosis in solid tumors, including small-cell lung cancer [59], renal cell carcinoma [60] and glioma [61], remaining a barrier for effective innate immune attack of the tumor cells. Antibody-mediated inhibition of CD47 has been shown to increase macrophage phagocytosis and polarization towards a tumor-attacking phenotype in liquid and solid tumors [62], increasing animal survival. Downregulation of CD47 in the GSC population, which can be achieved by inhibiting fibulin-3 as shown here, could prove significant as an approach complementary or alternative to direct CD47 blockade, in order to promote immune reactivation against the tumor.

Overall, the work presented here showcases fibulin-3, an ECM protein highly upregulated in malignant gliomas, as a widespread immunomodulatory component in the microenvironment of these tumors. Fibulins, neural chondroitin sulfate proteoglycans, hyaluronic acid, tenascins, fibronectin isoforms, osteopontin, and fibrillar collagens, have started to become recognized as bona-fide modulators of immune responses in glioma [63], either by direct signaling on immune cells, indirect signaling loops elicited in the tumor cells – as shown here for fibulin-3 –, or by changing the mechanical properties of the tissue scaffold. We acknowledge that we have not investigated here a possible direct effect of fibulin-3 on macrophages’ signaling or phenotype. This is an ongoing investigation in our laboratory, which requires separating any direct effects of fibulin-3 from the effects of the immunomodulatory signals upregulated in the tumor cells by fibulin-3. Nevertheless, our present results already emphasize this ECM protein as an actionable target that can be inhibited to reduce the expression of immunosuppressive signals in GBM cells, leaving them exposed to immune attack. We suggest that fibulin-3, as well as other pericellular ECM proteins, should be explored as targets to contribute to successful immunotherapy of aggressive brain tumors.

## Acknowledgements

This work was supported by research grants from the National Institutes of Health (1R21NS114615, MSV) and the US Department of Defense - PRCRP (CA-170319, MSV). We thank the charitable funding support provided by the SUNY Upstate Foundation - Debbie’s Brain Cancer Research Fund (to MSV)

