## Supplementary Materials for "Inhibition of the extracellular matrix protein fibulin-3 reduces immunosuppressive signaling in tumor stem cells and increases macrophage activation against glioblastoma"

Supplementary Tables I and II

Supplementary Figures S1 to S5

**Supplementary Table I:**

Antibodies used for Western blotting, flow cytometry, or immunohistochemistry

**Supplementary Table II:**

Sequences of oligonucleotides and primers used for q-RTPCR

**Supplementary Figure 1:** *Validation of humanized mAb428.2 equivalence to original mAb428 antibody.* Using the original $V_H$ and $V_L$ sequences of mouse mAb428.2 (Nandhu et al., Clin. Cancer Res. 2018), five humanized $V_H$ and $V_L$ variants were generated and cloned in a human IgG1 backbone (Absolute Antibody Ltd.), generating a total of 25 humanized mAb428.2 variants (*v1-v25*). All the variants were expressed in HEK293 cells, purified from culture medium to homogeneity, and tested against human fibulin-3 by indirect ELISA. **A)** Graph representing the production yield of mAb428.2 humanized variants (expressed as fold over the production of mouse chimeric mAb428.2 in HEK293 cells) and their binding affinity for fibulin-3. The red open dot represents the mouse monoclonal antibody. **B)** Percent of each humanized variant that remains as stable monomeric IgG in phosphate-buffered saline solution, compared against the mouse antibody source (*chimeric*). The images show non-reducing and reducing SDS-PAGE of purified mouse chimeric mA428.2 antibody (including a minor unstable product, arrow) and the humanized mAb428.2 chosen for this study. **C)** Representative Western blots of purified fibulin-3 (100 ng) detected with each humanized variant (tested at a concentration of 1 μg/mL). **D)** Representative ELISAs of the three humanized variants chosen for production (v5, v7, v10) because of their high production yield and high affinity against fibulin-3 compared to the original mouse antibody. The humanized variant hmAb428.2.v10 was used in the present study.

**Supplementary Figure 2:** *Quantitative analysis of TAM infiltration in intracranial tumors.* **A)** Comparison of intracranial tumor models with fibulin-3 knockdown versus their controls. **B)** Comparison of intracranial tumor models with fibulin-3 overexpression versus their controls. **C)** Comparison of an intracranial tumor model treated with anti-fibulin-3 mAb428.2 versus control IgG. For each tumor model, the results shown are as follows (left to right): 1) TAM infiltration ($IBA1^+$ area in the tumor); 2) IBA/CD206 co-expression in TAMs; 3) number of cells per mm2 (cell density control); and 4) area analyzed for each tumor section (tumor area control). Results for each parameter analyzed by Student's t-test with Welch's correction.

**Supplementary Figure 3:** *Correlation of fibulin-3 expression with immunosuppressive signals in GBM.* **A)** Correlation of fibulin-3 expression with IL-10 and the checkpoint genes CD80/CD86, observed in the TCGA GBM dataset. **B)** Correlation of fibulin-3 expression with an immunosuppressive signature (CSF-1, TGFβ, IL-10, CD47, CD274, CD86) observed in the CGGA GBM dataset. Data from the TCGA and CGGA datasets were recovered from the aggregator website Gliovis.

**Supplementary Figure 4:** *Anti-fibulin-3 treatment decreases immunosuppression in GBM.* Intracranial tumors (GBM09 model) treated locally with mAb428.2 (shown in **Figure 5**) were freshly resected and processed for qRT-PCR utilizing mouse-specific primers. Results show the comparative mRNA expression of several genes associated with TAM immunosuppression (CD206, ARG1, CD163), pro-inflammatory macrophage phenotype (TNF-α, IL-1β), and pro-tumoral phenotype observed in macrophages associated with rebound GBM (IGF-1, Quail et al., Science 2016). Results were compared by multiple paired t-tests with correction for multiple comparisons.

**Supplementary Figure 5:** *Anti-fibulin-3 triggers macrophage attack against syngeneic GBM cells*. fLuc-expressing GBM cells (GBM09, GBM34, and GL261) were co-cultured with a macrophage cell line (THP-1) or primary macrophages as indicated in the figure panels, in presence of anti-fibulin-3 mAb428.2 or its control IgG. The results show the extent of tumor cell death after 48 h of co-culture. Data were normalized to fLuc signal from cultures of GBM cells alone, treated with the same antibodies. Results were analyzed by Student's t-test for each GBM cell line. *PBMC:* macrophages derived from peripheral blood mononuclear cells *BMDM:* bone marrow-derived macrophages.

**Kundu et al., Figure S1**

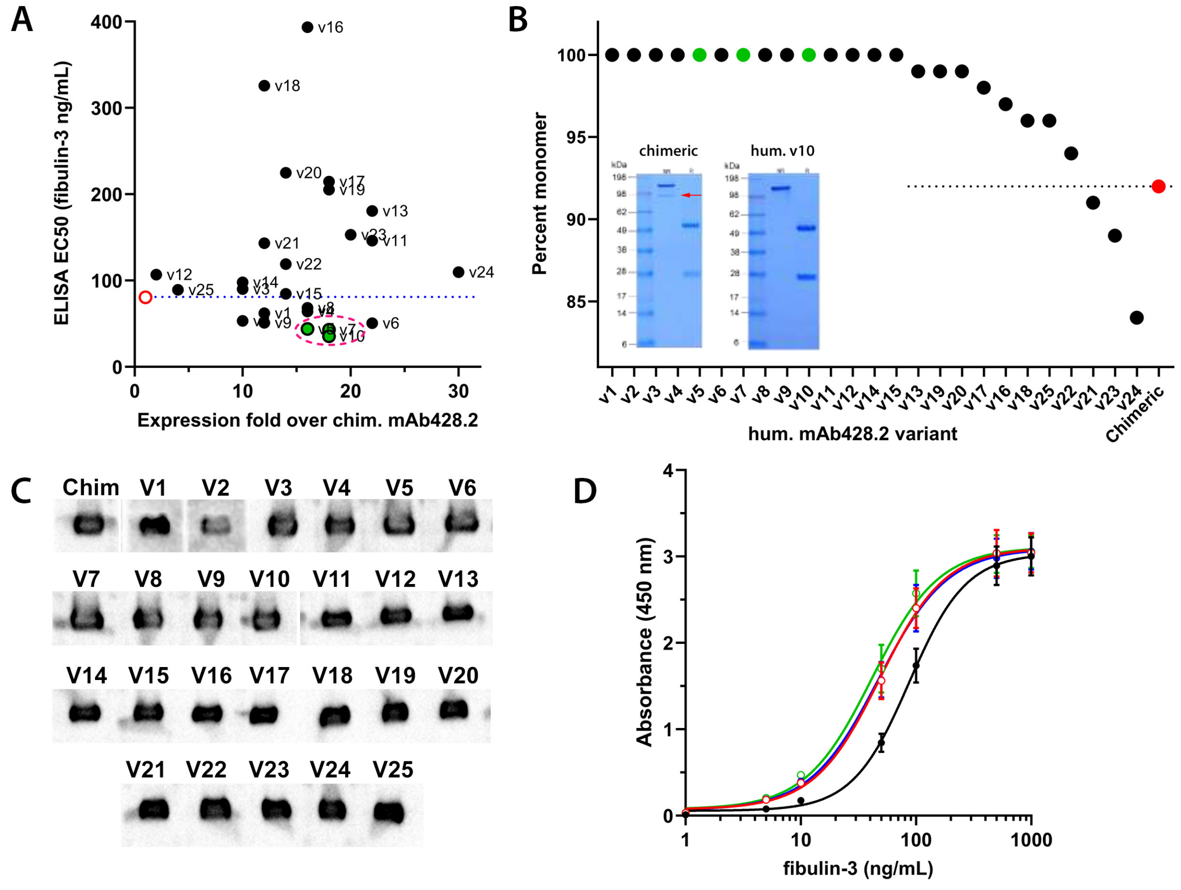


| | EC50 (ng/mL) | EC50 95% CI (ng/mL) |
|---|---|---|
| chim. mAb428.2 | 87.44 | 70.45 to 130.3 |
| hum mAb428.2 V5 | 48.57 | 31.74 to 66.11 |
| hum mAb428.2 V7 | 47.33 | 30.91 to 64.98 |
| hum mAb428.2 V10 | 41.54 | 25.94 to 58.41 |

Kundu et al., Figure S2

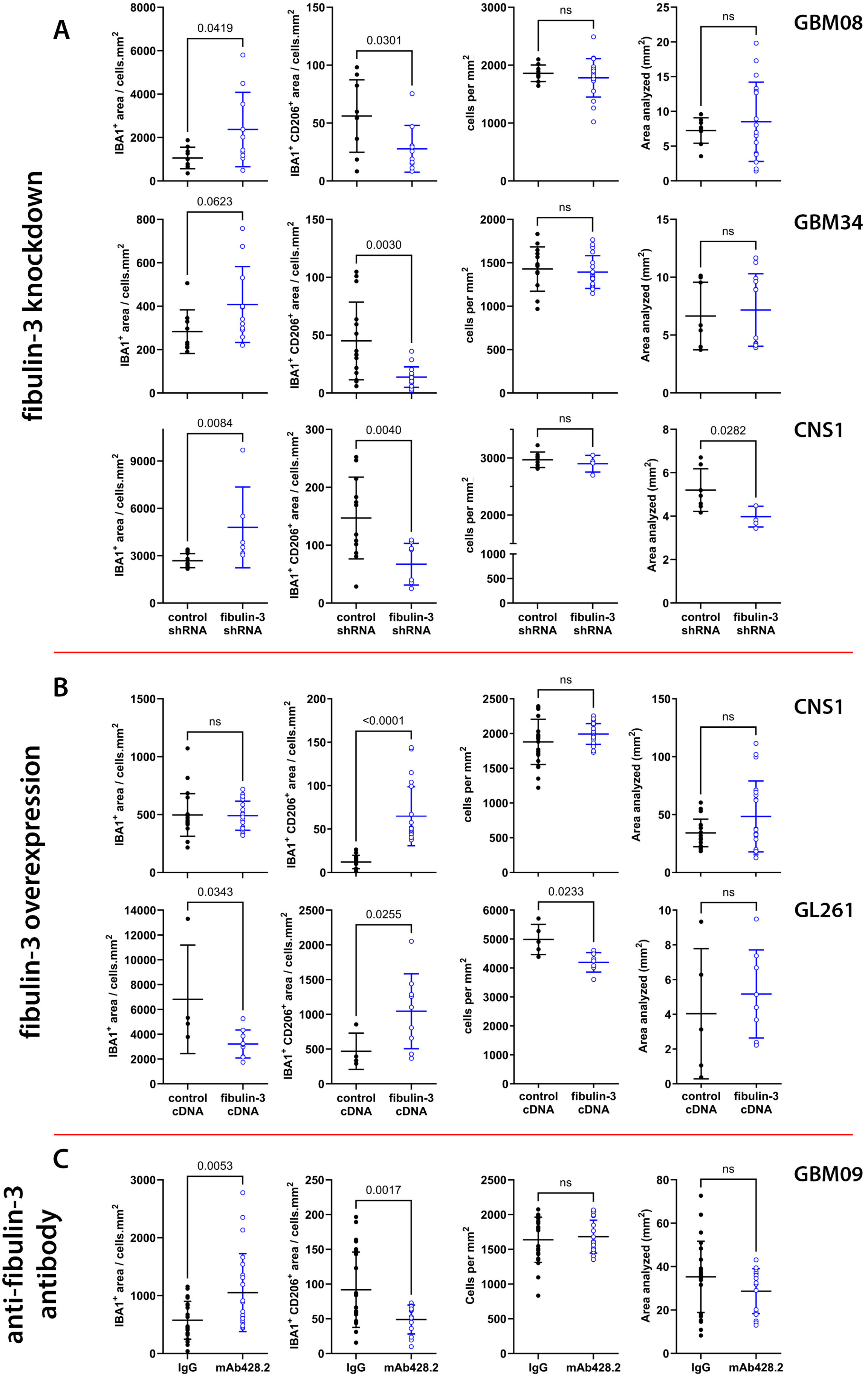

Kundu et al., Figure S3

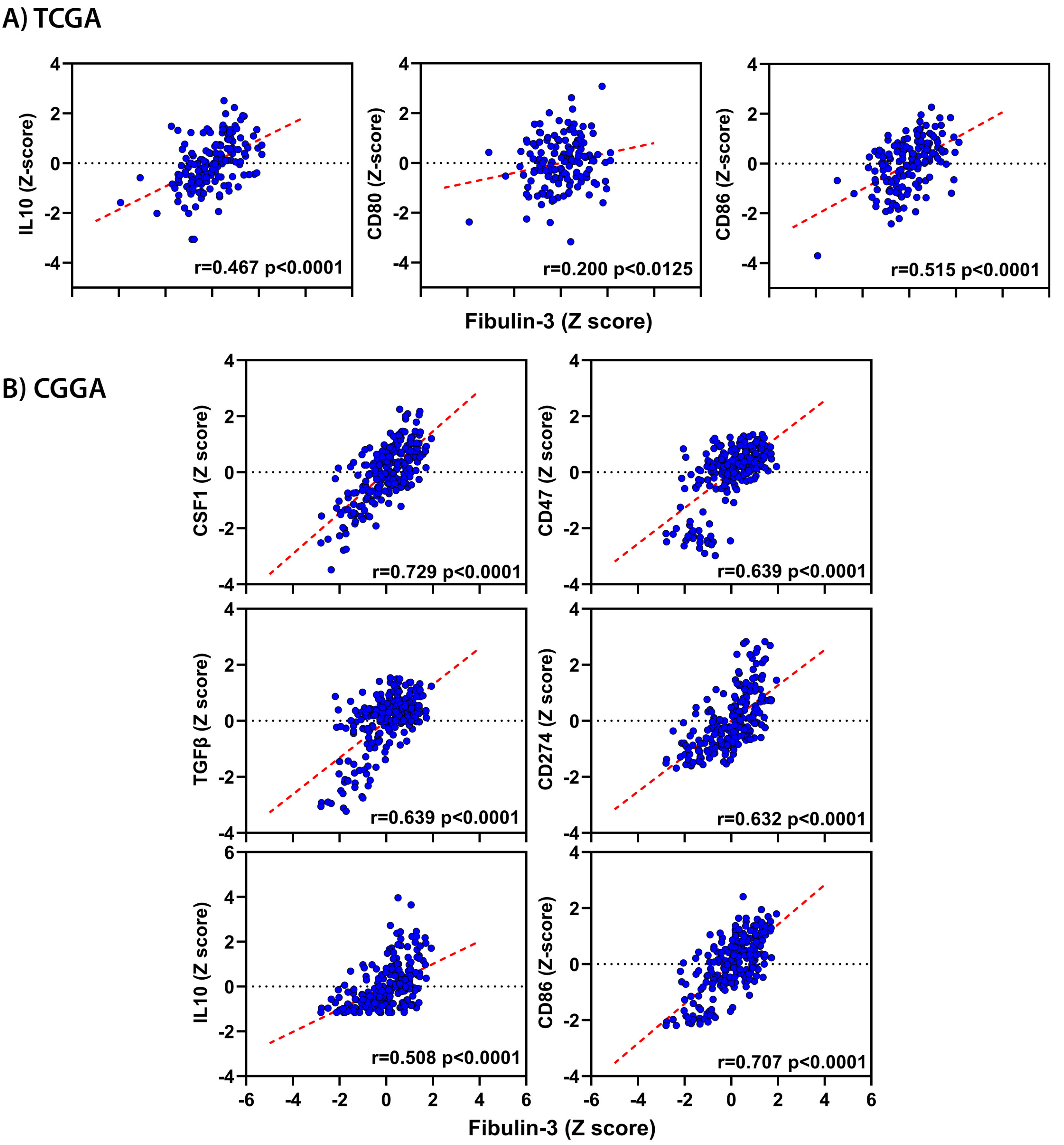

Kundu et al., Figure S4

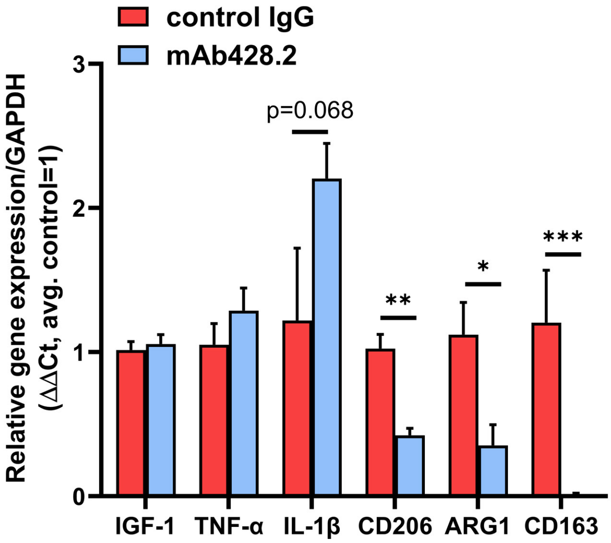

Kundu et al., Figure S5

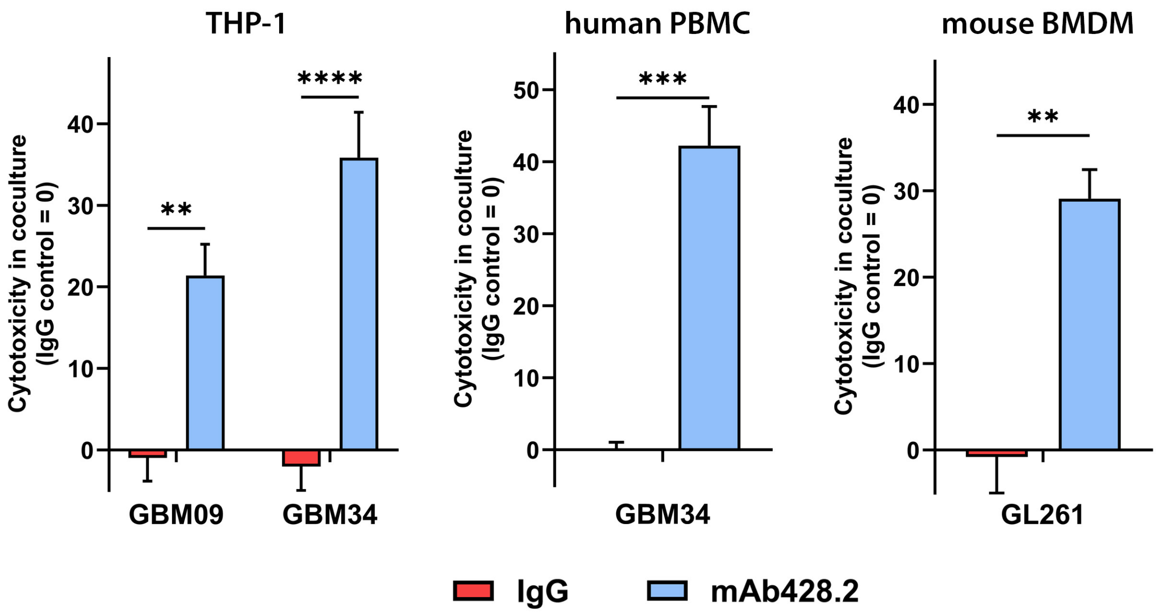

Kundu et al.

**Supplementary table I:**

**Antibodies used for Western blotting, flow cytometry, or immunohistochemisty**

| Antibody | Species | Source | Catalog # | Usage |
|---|---|---|---|---|
| CD11b conj. PE | Human | Miltenyi Biotec | #130-113-806 | Flow cytometry |
| CD206 conj. BV421 | Rat | BioLegend | #141717 | Flow cytometry |
| CD45 conj. APC | Rat | BioLegend | #103112 | Flow cytometry |
| CD45 conj. BV650 | Mouse | BioLegend | #304043 | Flow cytometry |
| Isotype control APC | Rat | Bio Legend | #400611 | Flow cytometry |
| Isotype control BV421 | Rat | Bio Legend | #400535 | Flow cytometry |
| Isotype control BV650 | Mouse | BioLegend | #400163 | Flow cytometry |
| isotype control PE | Human | Miltenyi Biotec | #130-113-450 | Flow cytometry |
| CD16/CD32 (non-specific Fc blocking) | Rat | BioLegend | #101302 | Flow cytometry |
| CD206 | Goat | R&D Systems | #AF2535 | IHC |
| IBA1 | Rabbit | Wako | #NB1001028SS | IHC |
| CD47 | Rabbit | Cell Signaling Technology | #6300 | Western blotting |
| CSF-1 | Mouse | Santa Cruz Biotechnology | sc-365779 | Western blotting |
| Fibulin-3 | Mouse | Santa Cruz Biotechnology | #sc-33722 | Western blotting |
| phospho NF-kB P65 | Rabbit | Cell Signaling Technology | #3033 | Western blotting |
| total NF-kB P65 | Rabbit | Cell Signaling Technology | #82425 | Western blotting |
| Vinculin | Mouse | R&D Systems | #MAB6896 | Western blotting |

**Kundu et al.**

**Supplementary table II:**

**Sequences of oligonucleotides and primers used for q-RTPCR**

| Gene | Oligo / Primer | Human-specific or multispecies sequence |
|---|---|---|
| Fibulin-3/EFEMP1 | siRNA1 | 5'- CACGCAATGCCACTGACGGATA |
| Fibulin-3/EFEMP1 | siRNA2 | 5'- CACAACGTGTGCCAAGACATA |
| P65/RelA | siRNA1 | 5'- AAGATCAATGGCTACACAGGA |
| P65/RelA | siRNA2 | 5'- CCGGATTGAGGAGAAACGTAA |
| ARG1 | Forward | 5'- AAGCAGACCAGCCTTTCTCA |
| ARG1 | Reverse | 5'- GCCAAGTCCAGAACCATAGG |
| CD163 | Forward | 5'- GGTGAATTTCTGCTCCATTCA |
| CD163 | Reverse | 5'- TGAGCCACACTGAAAAGGAA |
| CD206 | Forward | 5'- CTACAAGGGATCGGGTTTATGGA |
| CD206 | Reverse | 5'- TTGGCATTGCCTAGTAGCGTA |
| CD47 | Forward | 5'- ATGCATGGCCCTCTTCTGA |
| CD47 | Reverse | 5'- TTTGAATGCATTAAGGGGTTCCT |
| CD80 | Forward | 5'- GGCCCGAGTACAAGAACCG |
| CD80 | Reverse | 5'- TCGTATGTGCCCTCGTCAGAT |
| CSF-1 | Forward | 5'- CCAGCAACTGGAGAGGTGTC |
| CSF-1 | Reverse | 5'- GCAGCTGCAGGAACTCTCTT |
| Fibulin-3/EFEMP1 | Forward | 5'- TTTTGCTGTGCTGTGCAAGG |
| Fibulin-3/EFEMP1 | Reverse | 5'- CAGTGCATTGCGTGTACGTG |
| IL-10 | Forward | 5'- GCCACCCTGATGTCTCAGTT |
| IL-10 | Reverse | 5'- GTGGAGCAGGTGAAGAATGC |
| IL-1β | Forward | 5'- CCTCTCTCACCTCTCCTACTCA |
| IL-1β | Reverse | 5'- CTGCTACTTCTTGCCCCCTTT |
| INFγ | Forward | 5'- ACAGGGAAGCGAAAAAGGAGT |
| INFγ | Reverse | 5'- GCAGGCAGGACAACCATTAC |
| MIF | Forward | 5'- CGGACAGGGTCTACATCAACTA |
| MIF | Reverse | 5'- TCTTAGGCGAAGGTGGAGTT |
| P65/RelA | Forward | 5’- ACTGCCGAGCTCAAGATCTG |
| P65/RelA | Reverse | 5’- TCCCGTGAAATACACCTCAA |
| PD-L1 | Forward | 5'- TGGCATTTGCTGAACGCATTT |
| PD-L1 | Reverse | 5'- TGCAGCCAGGTCTAATTGTTTT |

| | | |
|---|---|---|
| TGFb | Forward | 5'- CAATTCCTGGCGATACCTCAG |
| TGFb | Reverse | 5'- GCACAACTCCGGTGACATCAA |
| TNF-α | Forward | 5'- CCCTGTGAGGAGGACGAACA |
| TNF-α | Reverse | 5'- TTTGAGCCAGAAGAGGTTGAGG |
| GAPDH | Forward | 5'- TTGCCCTCAACGACCACTTT |
| GAPDH | Reverse | 5'- TGGTCCAGGGGTCTTACTCC |
| 18S RNA | Forward | 5'- AACTTTCGATGGTAGTCGCCG |
| 18S RNA | Reverse | 5'- CCTTGGATGTGGTAGCCGTTT |
| **Gene** | **Primer** | **Mouse-specific Sequence** |
| ARG1 | Forward | 5'- TCACCTGAGCTTTGATGTCG |
| ARG1 | Reverse | 5'- CTGAAAGGAGCCCTGTCTTG |
| CD163 | Forward | 5’- TCTCCACACGTCCAGAACAG |
| CD163 | Reverse | 5’- CCTCGTCACCTTGGAAACAG |
| CD206 | Forward | 5'- AGTGGCAGGTGGCTTATG |
| CD206 | Reverse | 5'- GGTTCAGGAGTTGTTGTGG |
| Cd47 | Forward | 5’- AAATGGATAAGCGCGATGCC |
| Cd47 | Reverse | 5’- GGCTGATCCTTGGTCAGTGT |
| Csf-1 | Forward | 5’- GTGTCAGAACACTGTAGCCAC |
| Csf-1 | Reverse | 5’- TCAAAGGCAATCTGGCATGAAG |
| IGF1 | Forward | 5’- GCAACACTCATCCACAATGC |
| IGF1 | Reverse | 5’- AGCTGGACCAGAGACCCTTT |
| IL-1β | Forward | 5'- GCTTCAGGCAGGCAGTATC |
| IL-1β | Reverse | 5'- AGGATGGGCTCTTCTTCAAAG |
| TNF-α | Forward | 5'- CCACCACGCTCTTCTGTCTAC |
| TNF-α | Reverse | 5'- AGGGTCTGGGCCATAGAACT |
| GAPDH | Forward | 5'- GGTTGTCTCCTGCGACTTCA |
| GAPDH | Reverse | 5'- GCCTCTCTTGCTCAGTGTCC |
